# A Novel, Fast and Efficient Single-Sensor Automatic Sleep-Stage Classification Based on Complementary Cross-Frequency Coupling Estimates

**DOI:** 10.1101/160655

**Authors:** Stavros I. Dimitriadis, Christos Salis, David Linden

## Abstract

**Objective:** Limitations of the manual scoring of polysomnograms, which include data from electroencephalogram (EEG), electrooculogram (EOG), electrocardiogram (ECG) and electromyogram (EMG) channels, have long been recognised. Manual staging is resource-intensive and time-consuming and considerable efforts have to be spent to ensure inter-rater reliability. There is thus great interest in techniques based on signal processing and machine learning for a completely Automatic Sleep Stage Classification (ASSC).

**Methods:** In this paper, we present a single EEG-sensor ASSC technique based on dynamic reconfiguration of different aspects of cross-frequency coupling (CFC) estimated between predefined frequency pairs over 5s epoch lengths. The proposed analytic scheme is demonstrated using the PhysioNet Sleep European Data Format (EDF) Database using 20 healthy young adults with repeat recordings.

**Results:** We achieved very high classification sensitivity, specificity and accuracy of **96.2 ± 2.2%, 94.2 ± 2.3%**, and **94.4 ± 2.2% across 20 folds**, respectively and high mean F1-score (92%, range 90–94%) when multi-class Bayes Naive classifier was applied.

**Conclusions:** Our method outperformed the accuracy of previous studies on different datasets but also on the same database.

**Significance:** Single-sensor ASSC makes the whole methodology appropriate for longitudinal monitoring using wearable EEG in real world and lab-oriented environments.

## 1. Introduction

Sleep as a basic human function is characterized by continuous alterations in brain, muscle, eye, heart and respiratory activity. These multi-dimensional alterations are monitored with appropriate equipment in a sleep lab and measured during a full night of sleep. Typically, these polysomnographic recordings include the electroencephalogram (EEG), electro-oculogram (EOG), electromyogram (EMG) and electrocardiogram (ECG). Physiologically, sleep stages can be split into two types: rapid eye movement (REM sleep) and non-rapid eye movement (non-REM sleep) (Steriade and McCarley, 1990). The latter consists of 4 stages (called N1, N2, N3 and N4). The process of assigning to every epoch of polysomnographic recordings a sleep stage is called sleep scoring. Sleep staging is a very important step in sleep research and for the clinical interpretation of the polysomnogram (for a review see; Aboalayon et al., 2016).

In clinical daily routine, sleep studies are performed for the diagnosis of pathologies like circadian rhythm disorders, epilepsy, sleep apnea, insomnia and hypersomnia (Panossian and Avidan, 2009). Sleep scoring is often based on visual inspection of the polysomnographic recordings to establish a hypnogram that represent dynamically the different sleep stages. Sleep experts usually follow well established rules for manual sleep scoring based on state of the art guidelines (Hobson, 1969). Rechtschaffen and Kales (1968) introduced rules for the labelling of each segment of 30 s as Awake, S1–S4 or REM sleep stage. A more recent sleep manual proposed by the American Academy of Sleep Medicine (AASM) in 2007 (Iber et al., 2005), combines the non-REM stages S3 and S4 into a single stage of deep sleep (called N3), also known as slow-wave sleep (SWS). Both manuals suggest the use of EEG channels, two in the R&K (Hobson, 1969) manual and three in the AASM manual (Iber et al., 2005), 2 EOG electrodes and one EMG electrode.

Sleep stage scoring is the gold-standard for the analysis of human sleep (Agarwal and Gotman, 2005). While many sleep labs use the traditional manual sleep stage scoring of the neurophysiological recordings to produce the hypnogram, recent years have witnessed a burst of proposed methods for automatic or semi-automatic sleep staging (e.g. Agarwal and Gotman, 2005; Becq et al., 2005; Berthomier et al., 2007; Ma et al., 2011; Itil et al., 1969; Koley and Dey, 2012; Krakovská and Mezeiová, 2011; Larsen and Walter, 1970; Schaltenbrand et al., 1996; Sheng-Fu et al., 2012; Pan et al., 2012; Stanus et al., 1987; Gudmundsson et al., 2005; Šušmáková and Krakovská, 2008; Huang et al., 2002). The different methods for ASSC generally extracted features from the signals to analyze each temporal segment (epoch) and use classification algorithms to detect/predict the sleep stages (Güneş et al., 2010; Zoubek et al., 2007; Sousa et al., 2015). These features focused on time-domain analysis (Tsai et al., 2009; Gudmundsson et al., 2005; *Khalighi et al., 2011*), frequency-domain analysis (Zhovna and Shallom, 2008; Yu et al., 2012) and time-frequency-domain analysis (Ebrahimi et al., 2008; Li et al., 2009).

Complementarily, complexity and nonlinear measures have been successfully used (Kuo and Liang, 2011; Phan et al., 2013; Fell et al., 1996). In some ASSC systems, an appropriate preprocessing step of manipulating the selected features was added prior to the classification step. This preprocessing step includes feature selection and/or dimensionality reduction (Koley and Dey, 2012; Zoubek et al., 2007; Sen et al., 2014). The main scope of this step is to reduce the dimension of the estimated features. A wide range of machine learning-based classification methods such as Linear Discriminant Analysis (LDA) (Sousa et al., 2015; Weiss et al., 2011), Artificial Neural Networks (ANN) (Liu et al., 2010; Dursun et al., 2012), Support Vector Machine (SVM) (Huang et al., 2013; Brignol et al., 2012; Yu et al., 2012; Lainef et al., 2015), K-Nearest Neighbor (KNN) (Kuo and Liang, 2011; Liu et al., 2010), Decision Trees (DT) (Schaltenbrand et al., 1996; Pan et al., 2012) and SVMs-DT (Lan et al., 2015) have been adopted for sleep stage classification.

The clinical uptake of ASSC systems has been hindered by low accuracy, sensitivity and specificity. The classification accuracies varied among the ASSC methods reported in the literature, ranging from 70% to 94%, while the sensitivity and specificity remained lower than 90%. Some studies on ASSC systems have considered using a single EEG channel but many issues remained unsolved like the high similarity of the EEG characteristics between REM and S1 (Lan et al., 2015; Krakovská and Mezeiová, 2011;Estrada et al., 2004), which has lowered the overall classification performance.

Cortical excitability following rhythmic changes produces neuronal oscillations with different cell population size and therefore spatial scale (Fries,2005). It is well-known that low-frequency rhythms are established as a dominant coupling mode in distant neuronal interactions and long temporal windows. In the opposite, high-frequency rhythms are established as a dominant coupling mode in local neuronal interactions and short temporal windows (von Stein and Sarnthein, 2000). The continuous interactions between anatomical substrates/backbone and the cortical oscillatory patterns give the brain the flexibility to simultaneously work at multiple spatio-temporal scales (Buzsaki and Draguhn, 2004).

The distinct neuronal oscillations that travel with a dominant frequency rhythm are not independent and especially the lower frequencies can modulate the oscillations of higher frequency brain rhythms (Jensen and Colgin, 2007). The interactions between brain rhythms with a different frequency profile is called cross frequency coupling (CFC), which can be categorized as phase synchronization, amplitude co-modulation (correlation of the envelope: Bruns and Eckhorn,2004) and phase-amplitude coupling (PAC; Pittman-Polletta et al., 2014; Dimitriadis et al., 2015, 2016a, b).

Phase-amplitude coupling (PAC) is believed to be the substrate of neural coding and information exchange between micro and macro scales of the neural ensembles of the brain (von Stein and Sarnthein, 2000; Jensen and Colgin, 2007; Axmacher et al., 2010; Canolty and Knight, 2010). It seems that low-frequency oscillations regulate the information exchange between brain areas by modulating the excitability levels of local neural ensembles (von Stein and Sarnthein, 2000; Fries, 2005), while their phase affects high-frequency activity also on the level of individual neurons and their spiking rates (Canolty and Knight, 2010). PAC facilitates interactions between neuronal ensembles with similar phase and quantifies the strength of interaction between high-frequency bands with the phase of lower-frequency bands within low-frequency-dependent temporal windows (Allen et al., 2011). To quantify the PAC between brain signals of different frequency profile, we used three basic algorithms: a) the one based on iPLV (Dimitriadis et al., 2015, 2016), b) the one based on the mean vector length-MVL (Canolty et al., 2006) and c) the Modulation Index-MI (Tort et al., 2008).

Another type of CFC interactions between brain areas that oscillate on a different frequency rhythm is the correlation of the envelope of two brain signals that encapsulates brain activity of different spectrum profile (Bruns and Eckhorn,2004). It was found that inter-frequency coupling and specifically the correlation of the amplitude envelopes between low and high-frequency components is established between non-phase coupled patches in awake monkeys between areas located within their visual cortex (Bruns et al., 2001). It seems that envelope-to-signal correlation is a complementary type of CFC to PAC and reflects the transmission of temporally modulated brain activity from a source area to a target area in many conditions. Since the mechanism of neuronal exchange of information underlying the nature of this CFC metric is basic and the type of coupling in non-linear, this CFC type is of general interest for exploring the basic communication mechanisms in many situations.

The formation of new memories demands the coordination of neural activity across widespread brain regions. In both humans and animals, the hippocampus is believed to support the formation of new associative or contextually mediated memories (Clemens et al., 2009). During the consolidation of new memories on a system-level, mnemonic representations of items, thoughts, new faces etc initially reliant on the hippocampus and after are thought to travel to neocortical sites for more permanent storage. Sleep has this privilege role for facilitating this information transfer (Born and Wilhelm,2012). Mechanistically, consolidation processes have been proved to be rely on systematic interactions between the three basic neuronal oscillations that characterizing non–rapid eye movement (NREM) sleep, slow-oscillations, spindles and ripples (Staresina et al.,2015). The hierarchical role of these three components was revealed via phase-to-amplitude coupling based on mean vector length estimator. A recent study untangled hippocampo-cortical CFC as the basic mechanisms mediates memory consolidation during sleep (Maingret et al., 2016). They provided a clear link between sharp-wave ripples, δ waves and ripples. Logothetis et al., (2012) demonstrates the CFC coupling between hippocampo-cortical areas during a subcortical silence and off-line memory consolidation. Amiri et al., (2016) demonstrates an enhanced PAC in deep sleep and also in the onset zone of focal epilepsy.

Based on the aforementioned knowledge regarding CFC brain interactions, the multiplexity of brain interactions across different daily and lab-oriented tasks and our knowledge regarding sleep functionality in both humans and primates, we aimed on the present study to explore the effectiveness of different CFC estimates to the automatic sleep stage classification in normal human populations.

In this paper, we propose a fast and efficient single-sensor ASSC that achieves multi-class classification by combining estimation of cross-frequency coupling with a multi-class Bayesian Naïve classifier. First, we estimated the relative power on predefined frequencies extracted via wavelet analysis. Afterward, complementary cross-frequency coupling (CFC) estimators were adopted based on: a) phase-to-amplitude coupling (PAC) to quantify how the phase of the lower frequency brain rhythms modulates the amplitude of the higher oscillations, b) the correlation of the envelopes, c) the modulation index and d) a proposed complex version of modulation index. The whole approach was followed in a temporal segment (epoch) of 5 secs and within the two EEG bipolar sensors (FPz-Cz & Pz-Oz) and between every possible pair of the studying frequency bands. In previous applications PAC has shown promise as a biomarker for amnestic mild cognitive impairment subjects (Dimitriadis et al., 2015a), dyslexia (Dimitriadis et al., 2016), or mild traumatic brain injury (Antonakakis et al., 2016). We extracted the most important features using the infinite feature selection and fed them into a multi-class Bayesian Naïve classifier following a 20-fold cross-validation scheme. The whole approach was also validated in a second dataset.

In brief, the proposed ASSC methodology, which is described in more detail below, consists of four main steps: (i) estimation of CFC estimates per each cross-frequency pair in temporal segments of 5 s length in both EEG channels) and the relative power within each frequency, (ii) feature selection using sequential selection and cross-validation within the training dataset and (iii) classification using a multi-class Bayesian Naïve classifier based on the features selected in (ii). External-validation of the proposed methodology in a second dataset using the features selected from the first one (iv).

The layout of the paper is as follows. In Section 2, we describe the subject population, the experiments that were performed, and the methods used for data pre-processing steps of the proposed pipeline and the classification procedure. Results are presented in Section 3, and Section 4 is devoted to the discussion.

## 2. Materials and Methods

The proposed sleep stage classification approach can be divided in four steps. Fig.1 demonstrates those four steps which can be summarized as follow:

1. Signal processing and extraction of wavelet components within the predefined frequencies
2. Estimation of the features based on relative power, different CFC estimators per frequency pair
3. Feature selection and finally
4. Classification of the sleep stages

**Figure 1.**
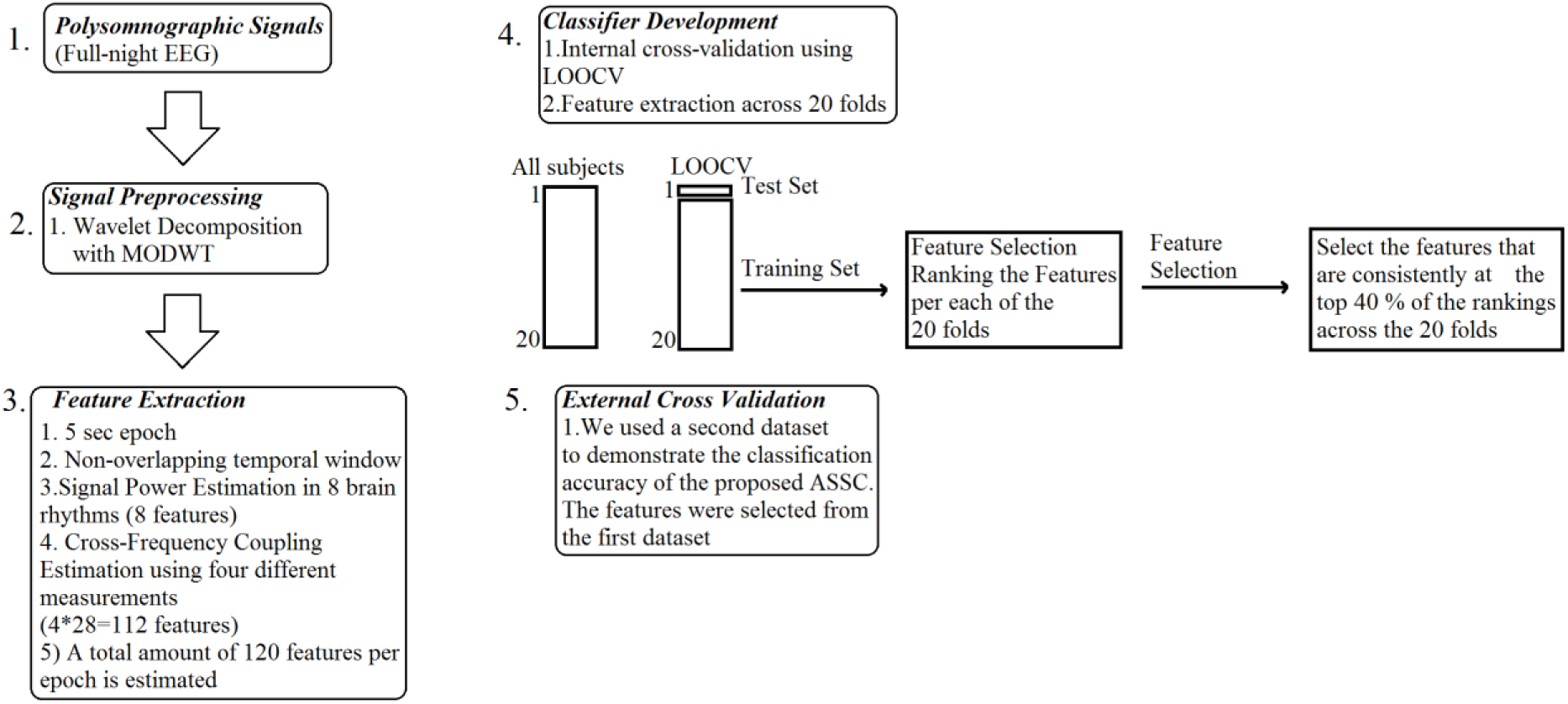
The outline of the methodology.

### 2.1. Polysomnographic database

The **polysomnographic** dataset that we used to present and evaluate the proposed novel methodology is a publicly available sleep PSG dataset (Kemp et al., 2000) demonstrated as part of the PhysioNet repository (Goldberger et al., 2000) that can be downloaded from (The Sleep-EDF Database [Expanded]”, Physionet.org). The brain activity was recorded from two electrodes-pairs, the Fpz-Cz and Pz-Oz, instead of the standard C3-A2 and C4-A1. The sleep stages were assigned to the following stages/conditions: wake (W), REM (R), non-R stages 1–4 (N1, N2, N3, N4), movement and not scored. The sleep scoring of each epoch of length 30s was realized by six experts following the Rechtschaffen and Kales guidelines (Hobson,1969). For our study, we removed from further analysis the very small number of movement and not scored epochs (not scored epochs were mostly at the beginning or end of each recording). We also keep N3 and N4 as distinct sleep stages. In the whole dataset, we detected 61 epochs with movements while only 17 had artifacts due to movements. Across the cohort, the maximum number of movement epochs was 12.

The adopted open-access dataset consists of 20 healthy subjects (10 male / 10 female), aged 25–34 years. There are two 20-h recordings per subject. EEG recordings were sampled at 100 Hz while epoch duration is 30 s.

### 2.2. Feature Extraction

Both EEG channel recordings were decomposed every 5s with Maximal Overlap Discrete Wavelet Tansform (MODWT) wavelet method and Daubechies wavelet filters (dau4). The wavelet signals were then mapped to one of the eight predefined frequency ranges. The reason why we preferred a wavelet decomposition over predefined bandpass filtering of the sleep EEG recordings is to get a more accurate temporal resolution of the well-established frequency ranges and to distinguish true from artefactual sleep activity. For that reason, we combined EOG and EMG recordings with wavelet time series to remove signals directly linked to eye movements and muscle activity. The predefined frequencies were: low-δ {0.1-1.5 Hz}, high-δ (K-Complex) {1.6-4 Hz}, θ {4-8 Hz},α_1_ {8-10 Hz}, α_2_ {10-13 Hz},β_1_ (spindle) {14-20 Hz},β_2_ {21-30 Hz} and γ_1_ {31 – 45 Hz}.

We splitted δ frequency band in order to capture slow wave activity and also to focus on well-known frequency profile of K-complex that suppresses cortical arousal in response to any external and second, plays a key role to sleep-based memory consolidation (Cash et al., 2009). Additionally, β brain rhythm was further splitted into low and high in order to capture sleep spindles activity via CFC. Sleep spidles occur during stage 2 sleep and are often follow the occurrence of K-Complex. Sleep spindles result from thalamo-cortical interactions and has been found: a) to suppress the presence of disruptive external sounds, a correlation has been revealed between the brain activity in the thalamus and the subject’s ability to be tranquile (Thanh Dang-Vu et al., 2010) and b) to be associated with the integration of new information into existing knowledge (Tamminen et al., 2010) as well as directed remembering and forgetting (Saletin et al., 2011; fast sleep spindles).

We estimated five types of features: 1) the relative signal power for each frequency band in the time domain based on the Maximal overlap discrete wavelet transform (MODWT), 2) the phase-to-amplitude coupling (PAC) that is estimated between every possible pair of frequencies, 3) the correlation coefficient between the envelopes of the frequency bands as an amplitude-amplitude cross-frequency coupling (AAC) estimator, 4) a novel complexed version of the modulation index (CMI) for estimating the phase-to-amplitude coupling between every possible pair of frequencies and 5) the original modulation index (MI). All the cross-frequency coupling estimators were computed between all possible pairs of the eight predefined frequency bands (8*7/2=28 combinations).

All features were estimated within each epoch of 30 s length and also by adopting a sliding non-overlapped window of duration of 5s, which resulted in 6 temporal non-overlapped segments per 30 s. The whole analysis leads to the extraction of 120 (8 relative signal power + 28 Phase-to-amplitude coupling (PAC) + 28 Amplitude-to-Amplitude coupling (AAC) + 28 Phase-to-Amplitude coupling (CMI) + 28 Phase-to-Amplitude coupling (MI)) total features per epoch. All the features were mapped in the [0,1] interval independently for each EEG channel.

#### 2.2.1 Relative Signal Power

EEG recordings of every 5s sub-epoch was decomposed using the Maximal overlap discrete wavelet transform (MODWT) wavelet method and Daubechies wavelet filters (dau4). The outcome of this process were time series with frequency profile that corresponds to the predefined frequency bands.

We estimated the relative power of each band-pass frequency signal segment in the time-domain with the following equations:

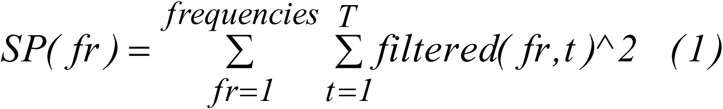

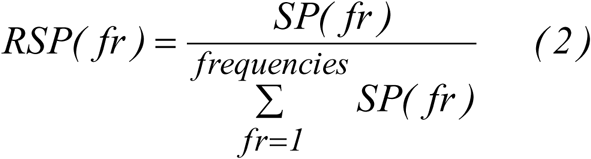

The first equation quantifies the signal power (SP) of each frequency as the sum of the filtered signal squared per sample (1) while equation (2) divides the SP by the sum of the SP from all the frequencies which gives the relative signal power (RSP). The whole approach was repeated for every epoch, sub-epochs, EEG sensor-pairs and subject.

#### 2.2.2 Cross-Frequency Coupling Estimations

##### 2.2.2.1 Phase-to-Amplitude Coupling Cross-Frequency Coupling (PAC)

CFC quantifies the strength of interactions between time series of different frequency content. It can be estimated both within and also between sensors (Canolty and Knight, 2010; Buzsáki, 2010; Buzsáki et al., 2013). CFC can be estimated between power – power, amplitude – amplitude and amplitude-phase representations of two time series with different frequency content. These representations can be derived by filtering twice one (within) or once two-time series (between). The most common type of CFC interaction is phase-to-amplitude coupling (PAC) and it is the most common in the literature (Voytec et al., 2010). The PAC algorithm for a single EEG sensor is described below.

Let x(i_sensor_, t), be the EEG time series at the i_sensor_-th recording site, and t=1, 2,.… T the sample points. Given a band-passed filtered signal x(i_sensor_,t), CFC is quantified under the notion that the phase of the lower frequency (LF) oscillations modulate the amplitude of the higher frequency (HF) oscillations. The following equations described the complex representations of both (LF) z_LF_(t) and (HF) oscillations z_HF_(t) produced via the Hilbert transform (HT[.]).

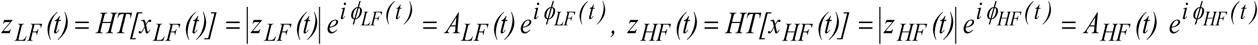

The next step of the PAC algorithm is the estimation of the envelope of the HF oscillation AHF(t) which then is decomposed via the wavelet and we selected the component within the frequency range of LF oscillations. Afterward, the resulting time series is again Hilbert transformed in order to get its phase time series that described phase dynamics φ’(t)

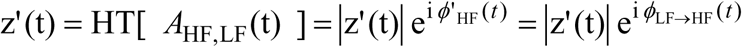

The aforementioned complex equation describes analytically the modulation of the amplitude of HF oscillation by the phase of LF oscillation.

The phase consistency between those two time series can be measured by the original phase locking value (PLV) estimator (Lachaux et al., 1999) but also from its imaginary portion of PLV. Imaginary part of PLV (iPLV) can be used as an synchronization index that quantifies the strength of CFC-PAC coupling.

PLV is defined as follows:

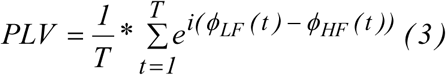

and the iPLV as follows:

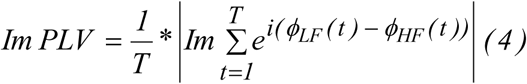

The iPLV is an estimator that is less affected compared to PLV from the volume conduction effect. Using iPLV for quantifying the strength of CFC interactions is an advantage over volume conduction. iPLV is more sensitive to non-zero phase lag and for that reason is more resistant to any self-interactions that are directly linked to volume conductions (Nolte et al., 2004). For further details and applications, an interested reader can read our previous work (Dimitriadis et al., 2016; Antonakakis et al., 2016).

In the present study, as was already mentioned we used the wavelet signals that correspond to the eight frequency bands which means that PAC is estimated for 8*7/2=28 cross-frequency pairs e.g. δ^φ^ − θ^A^,δ^φ^ − α1^A^ where φ and A denote the phase and amplitude of each frequency band. Figure 2 demonstrates the pre-processing steps of the PAC estimator using a 30s epoch from wake stage of subject 1.

**Figure 2.**
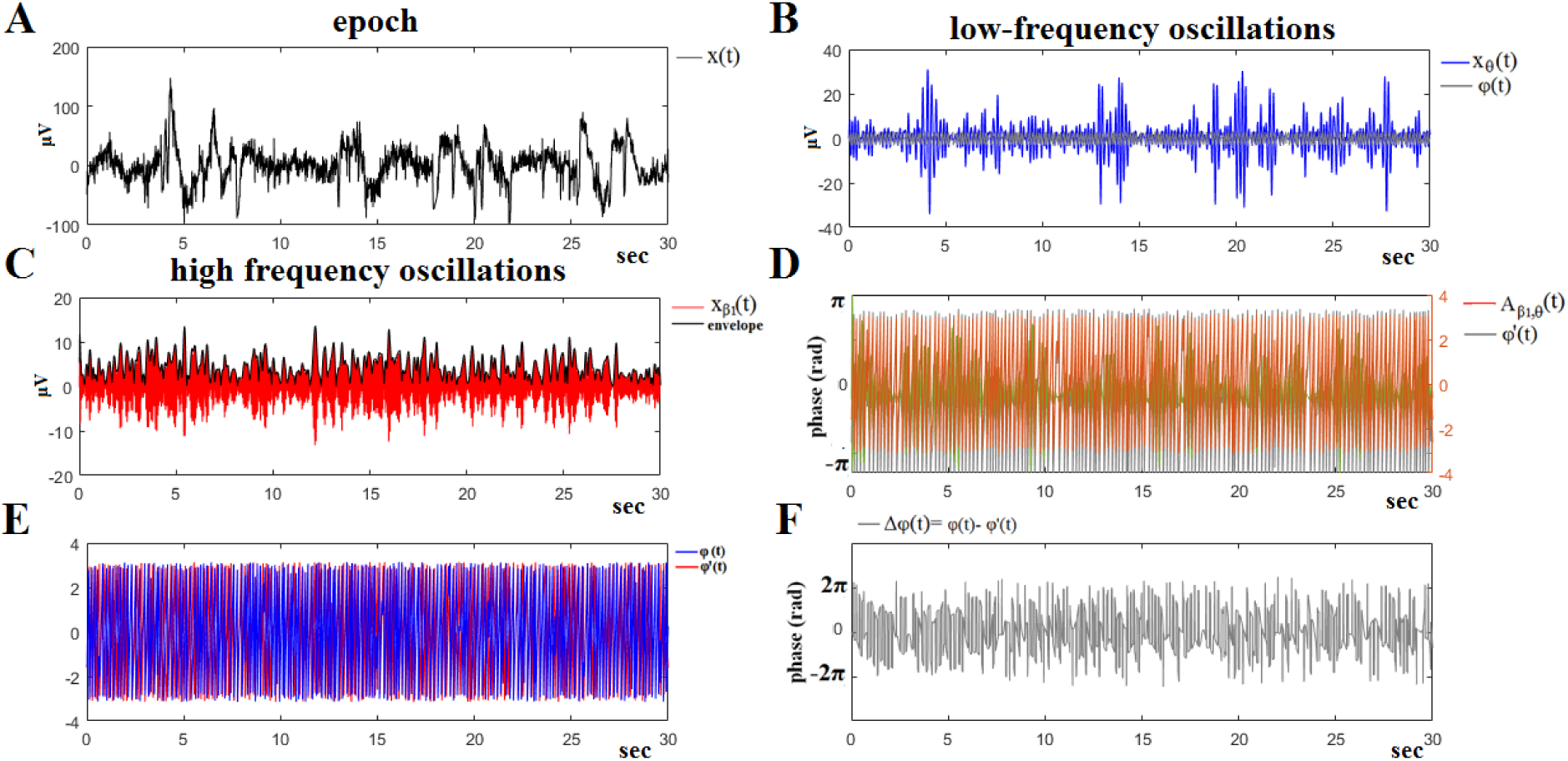
The algorithmic steps for PAC estimation. Using the first epoch EEG signal from Fpz-Cz sensor of subject 1 during wake condition **(A)**, from the P300 of an able subject (subject 6), we demonstrated the phase-to-amplitude coupling between θ and β_1_ rhythm. Firstly, the raw time series was band-pass filtered using a zero-phase order filter into **(B)** low-frequency θ (4– 8 Hz) component where its envelope (Hilbert transform) is extracted and into **(C)** a high-frequency β_1_ (13–20 Hz) component where via Hilbert transform its phase dynamics is estimated. **(D)** We then estimated both the amplitude and also the instantaneous phase of the band-passed β_1_ (13–20 Hz) component and we filtered the amplitude of this time series within the θ frequency range (4–8 Hz). This algorithmic step will give us the θ modulation within the lower β amplitude. **(E)** Afterward, we Hilbert transformed both the θ-filtered signal and the θ-filtered within the lower-β amplitude extracting the related phase dynamics and finally their phase consistency with iPLV. The phase differences of those two phase time series which is illustrated in **(F)**, will be the input in the iPLV estimator in order to quantify the strength of PAC coupling between θ and β_1_ rhythm and how the phase of the lower frequency component modulates the amplitude of the high amplitude

##### 2.2.2.2 Amplitude-to-Amplitude Cross-Frequency Coupling (AAC)

The complex analytic representations of each signal as derived via the MODWT approach zF(t) was then Hilbert transformed (HT[.]).

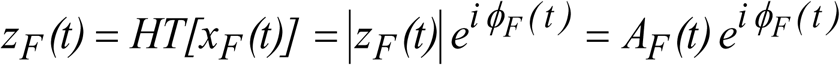

Next, the envelope of the frequency oscillations AF(t) was squared to express the power of the signal in the time-domain. Afterward, the correlation coefficient between every pair of the derived time series was estimated to express the associations between specific frequency pairs.

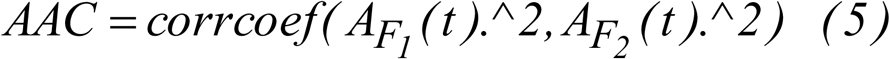

In the present study, as was already mentioned we used the wavelet signals that correspond to the eight frequency bands which means that AAC is estimated for 8*7/2=28 cross-frequency pairs e.g. δ^A^ - θ^A^,δ^A^ - α_1_^A^ where A denote the amplitude of the envelope of each frequency band. Figure 3 demonstrates the pre-processing steps of the AAC estimator using a 30s epoch from wake stage of subject 1.

**Figure 3.**
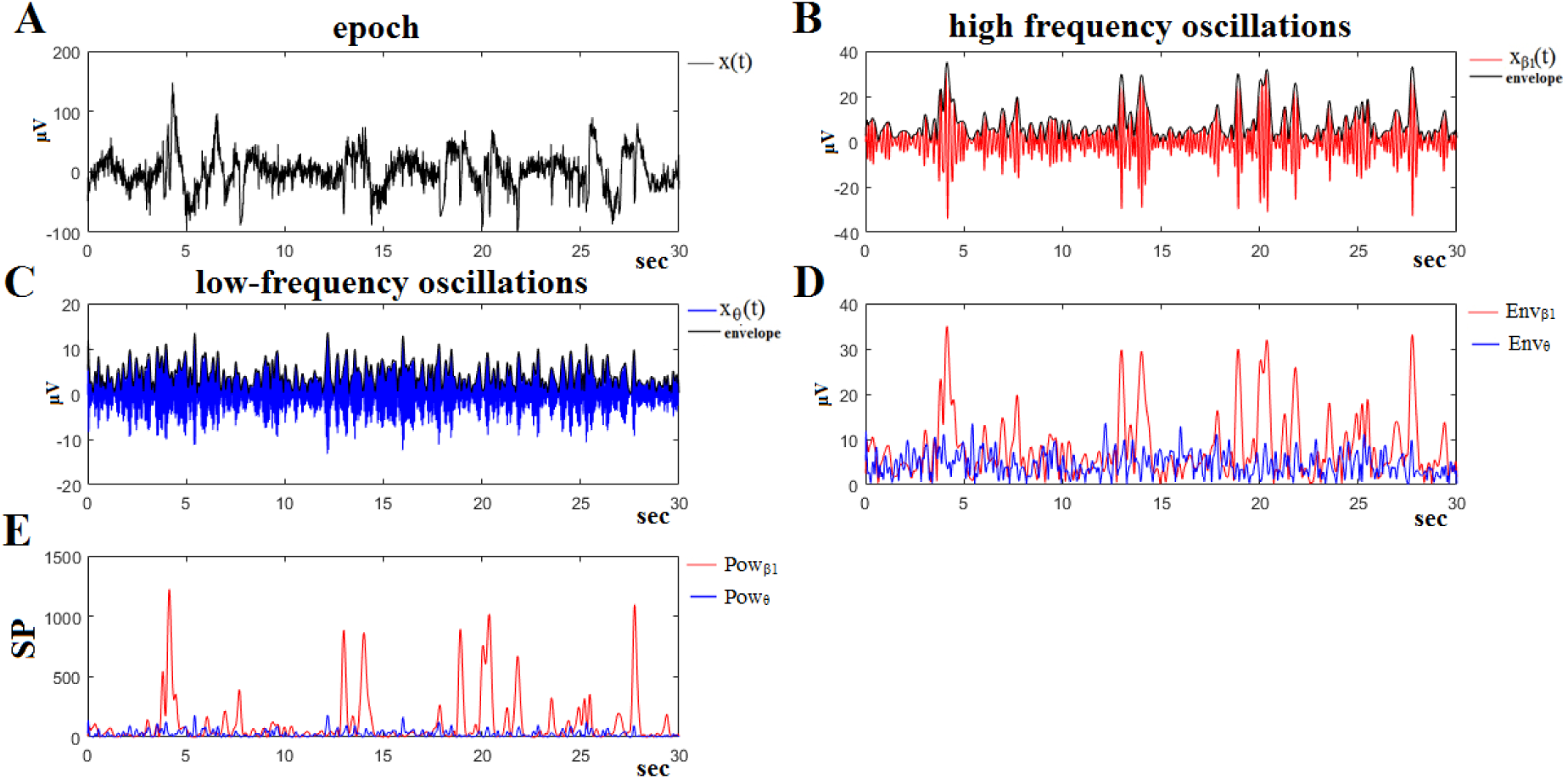
The algorithmic steps for AAC estimation. Using the first epoch EEG signal from Fpz-Cz sensor of subject 1 during wake condition **(A)**, we demonstrate the detection of coupling between θ and β_1_ rhythm. To estimate θ-β_1_ AAC, the raw signal was band-pass filtered into both **(B)** a high-frequency β_1_ (13–20 Hz) component where its instantaneous phase is extracted and a **(C)** low-frequency θ (4–8 Hz) component where its envelope is extracted as well as. **(D)** We then presented the envelopes of the band-passed θ (4–8 Hz) and β_1_ (13–20 Hz) into a common plot. **(E)** Finally, we estimated the point-wise squared time series of the envelopes to get the power time series. The AAC is estimated on these power time series using correlation coefficient.

##### 2.2.2.3 A Complex Version of the Modulation Index (CMI)

Modulation Index (MI) has been presented as a novel estimator for constructing a phase-amplitude plot (comodulogram) that demonstrates the strength of how the phase of a low-frequency modulates the amplitude of the high-frequency within a raw signal (Tort et al., 2010).

Let’s denote by *x*_*raw*_(*t*) the raw signal here one of the EEG recordings. The MI is calculated for every pair of frequencies creating a phase-amplitude plot. The steps for estimating the modulation index are the following:

1. First, *x*_*raw*_(*t*) is filtered at the two frequency ranges, the low-frequency *f*_*p*_ and the high-frequency *f*_*A*_. We denote the filtered signals as *x*_*fp*_(*t*) and *x*_*fA*_(*t*).
2. The time series of the phases of *x_fp_*(*t*) [denoted as ϕ*_fp_*(*t*)] is obtained from the standard Hilbert transform of *x_fp_*(*t*). The Hilbert transform is also applied to *x_fA_*(*t*) to extract the time series of the amplitude envelope of *x_fA_*(*t*) [denoted as *A_fA_*(*t*)]. A composite time series is constructed [ϕ*_fp_*(*t*), *A_fA_*(*t*)] giving the amplitude of the *fA* oscillation at each phase of the *fp* rhythm.
3. Next, the phases ϕ*_fp_*(*t*) are binned (here we used 20 bins of the 360^o^ of range 18^o^) and the mean of *A_fA_* over each phase bin is calculated. We denote by (*j*) the mean *A_fA_* value at the phase bin *j*.
4. At the last step, we normalize the mean amplitude by dividing each bin value by the sum over the bins

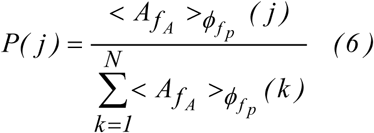

where *N* denotes the number of phase bins (here N=20s).

Here, instead of equation 6, we used the following transformation based on the composite signal

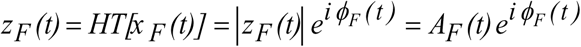

And is defined by the following equation.

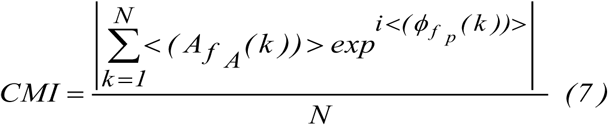

where we get the product of the mean *A_fA_* within each phase bin with the mean ϕ*_fp_* within each phase bin N.

In the present study, we employed the wavelet signals that correspond to the eight frequency bands which means that CMI is estimated each one for 8*7/2=28 cross-frequency pairs e.g. δ^A^ - θ^A^,δ^A^ - α_1_^A^ where A denote the amplitude of the envelope of each frequency band. Figure 4 demonstrates the pre-processing steps of the CMI estimator using a 30s epoch from wake stage of subject 1.

**Figure 4.**
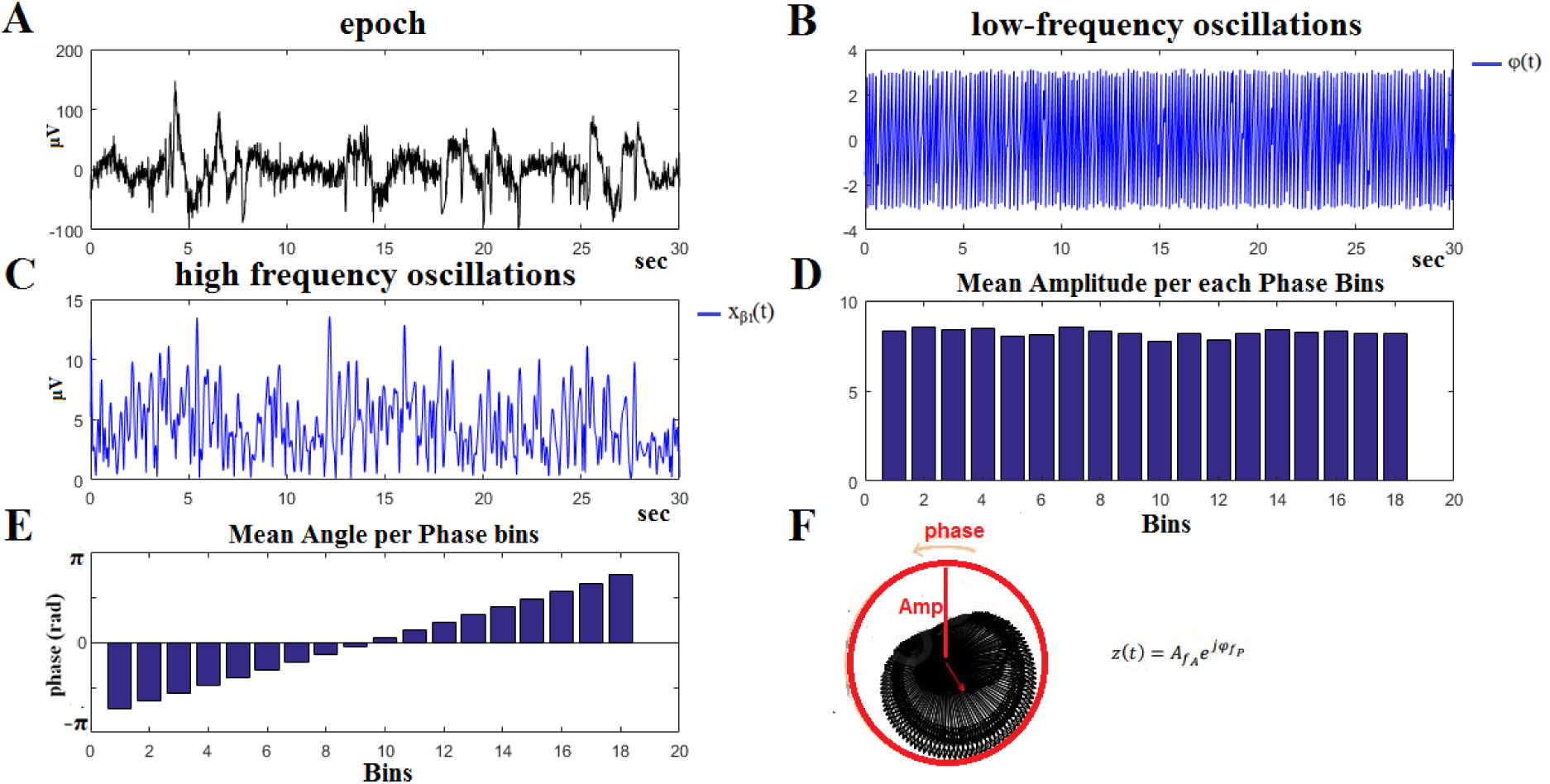
The algorithmic steps for CMI estimation. Using the first epoch EEG signal from Fpz-Cz sensor of subject 1 during wake condition **(A)**, we demonstrate the detection of coupling between θ and β_1_ rhythm. To estimate θ-β_1_ CMI, the raw signal was band-pass filtered into a **(B)** low-frequency θ (4–8 Hz) component where its phase is extracted and into **(C)** a high-frequency β_1_ (13–20 Hz) component where its envelope is extracted. **(D)** The mean amplitude of the high-frequency was estimated within each (E) phase bin extracted from the phase of the low-frequency. The CMI is estimated on based on the average complex valued of mean amplitude and mean phase within each bin. Here, we used 20 phase bins.

##### 2.2.2.4 The Modulation Index (MI)

The original defined MI (Tort et al., 2010) was estimated via the equation 6. Figure 4 demonstrates the pre-processing steps of the MI estimator using a 30s epoch from wake stage of subject 1. Equation 4 is calculated based on the amplitude and phase bins presented in Fig.4D,E.

Here, we used the wavelet signals that correspond to the eight frequency bands which means that MI is estimated for 8*7/2=28 cross-frequency pairs e.g. δ^A^ - θ^A^, δ^A^ - α_1_^A^ where A denote the amplitude of the envelope of each frequency band.

##### 2.2.2.5 A graphical Visualization of Cross-Frequency Interactions

For every epoch of 5 sec and for the four cross-frequency coupling estimators, we quantified the cross-frequency interactions. The possible pair-wise combinations of the eight brain rhythms are 28 while γ_1_ cannot be a modulator but only a modulated frequency. We visualized these 28 cross-frequency interactions using a graphical representation where the nodes denote the seven out of eight brain rhythms, the direction of arrows illustrate the direction of the modulation while the color encodes the strength of the coupling. In our example, we adopted epoch 1 from N1 of the first subject from the training dataset using phase-to-amplitude estimator. It is important to underline that in the present study, we used a single-sensor for the estimation of the four cross-frequency interactions and relative power estimates. For every epoch of 5 sec and for each of the four CFC estimates, we quantified the pair-wise cross-frequency interactions among the eight brain rhythms. A complementary visualization scheme for the cross-frequency coupling interactions is given in Fig.5 called comodulogram in a graphical layout compared to a tabular representation (Dimitriadis et al., 2015a, 2016a).

**Figure 5.**
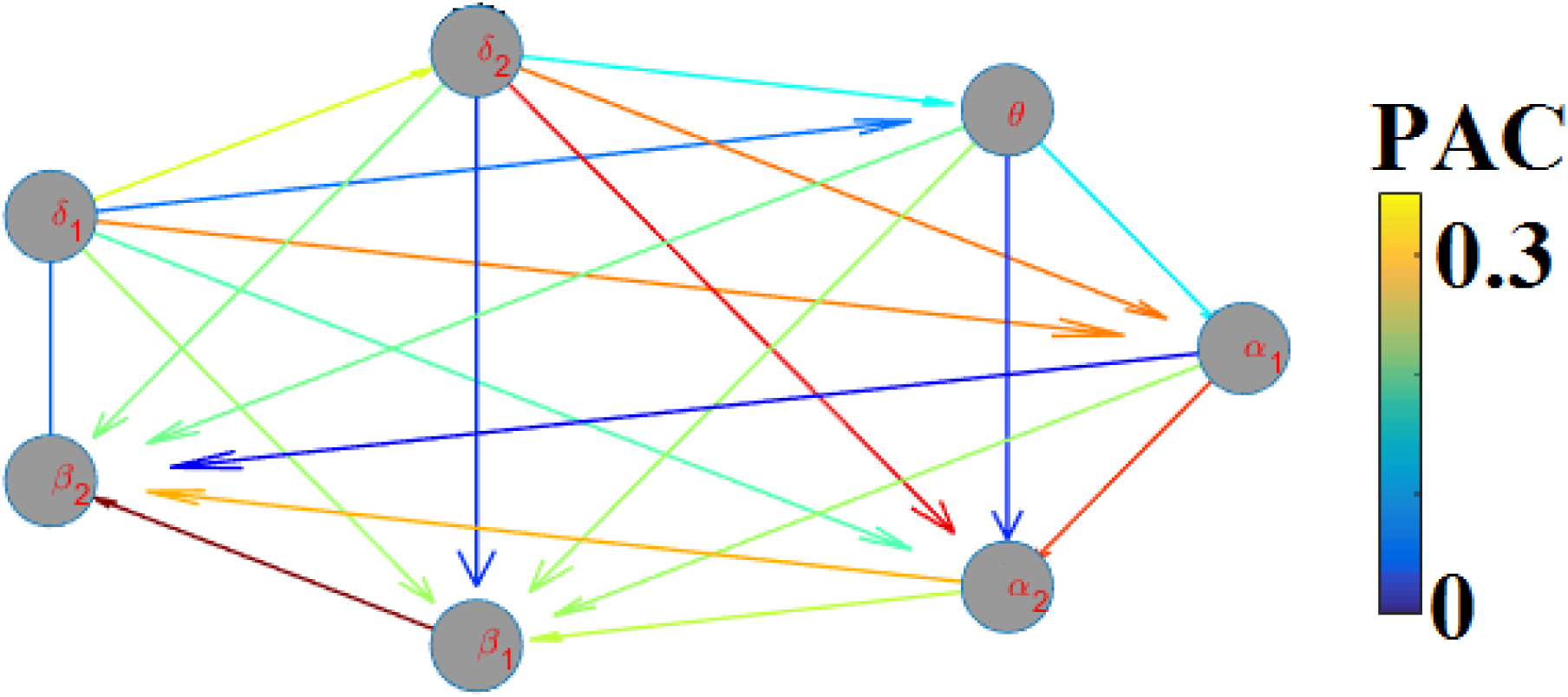
A graphical representation of the PAC-couplings for the first epoch of N1 sleep stage of subject 1 in the first dataset. Nodes represent the modulating brain rhythms, the arrows are directed to the modulated frequencies while the color refers to the strength of PAC. At each epoch of 5 sec and for each of the four cross-frequency coupling estimates, all possible pair-wise frequency interactions were estimated as they are demonstrated with the graphical representation.

### 2.3 Feature Selection

We ranked the whole set of estimated features using the infinite feature selection method introduced in (Deng et al., 2010). We applied the feature selection strategy to the training set in every fold of the leave-one out cross-validation scheme (LOOCV; see next section). Finally, we selected the features that were consistent selected in the top 40 % across the 20 folds. For every fold, we got a ranking of the 120 features and finally we selected the number of features that were consistently appeared on the top 40% across the 20 folds (see also in Fig.1).

### 2.4 Machine learning and classification

We accessed the generalizability of the whole approach by using 20-fold cross-validation. Specifically, in each fold we used the features extracted from a single subject for testing and all other recordings for training. The methodology was tested with both available EEG electrodes (Fpz-Cz and Pz-Oz). Each subject’s recordings were used only once for testing, thus obtaining a one-to-one correspondence of cross-validation folds and test subjects.

We report the scoring performance using the best electrode, which was Fpz-Cz. Group-averaged confusion matrices are reported to present the average classification performance per sleep stage while mean accuracy, sensitivity, specificity and F1-score are also reported.

We used five different classifiers: a) the k-nearest neighbour, b) the Bayesian Naïve multi-class classifier (Linear and Kernel), c) the extreme-learning machine (ELM) (Linear, RBF) (Huang et al., 2012), d) linear discriminant analysis (LDA) and e) the multi-class support vector machines (multi-SVM – libsvm toolbox) (Linear,RBF) (Chang and Lin, 2011).

To avoid the effect of imbalanced sleep stage representation and present true performance measures (accuracy, sensitivity, specificity, F1 score) and confusion matrices, we randomly sampling the sleep stages across each fold such as to get equal representation of each sleep stage across sleep stages and subjects. We repeated the 20-fold cross-validation 100 times.

### 2.5 Second Polysomnographic Database

The adopted second open-access dataset consists of 77 healthy subjects (36 male age *59.01*±*23.31* / 41 female *58.53*±*21.6*), aged 26–101 years. There are whole-night polysmnographic sleep recordings containing EEG (from Fpz-Cz and Pz-Oz electrode locations), EOG (horizontal), submental chin EMG, one from each subjects. EEG recordings were sampled at 100 Hz while epoch duration is 30 s.

### 2.6 Computational time of the proposed ASSC

Automatic sleep scoring by a neurologist demands 3-4 hours for a single hypnogram (Lainscsek et al., 2013). Our algorithm can estimate the selected features and produce the ASSC of a single subject based on the training set on averaged *8.9*±*1.3* mins (depending on the duration of the sleep recordings) using a multi-core processor and parallelized the MATLAB code in a MATLAB environment. We estimated the computational time on the second dataset which was *9.2*±*1.5*.

## 3. Results

### 3.1 Discriminative Features of Sleep Stages

In the following tables, we demonstrated the selected features from the pool of four complementary cross-frequency estimators (Table 1-4) adopted here and the relative power estimates (Table 5). We underlined with a ‘tick’ in Tables 1-5, the selected features which were 36 out of 120. Using a criterion of consistency of features on the top 40% (0.4*120=48 features) across the 20-folds, we finally selected 36 features from the first training set. The same set of features was used for an external validation of the whole approach to the second dataset.

**Table 1.** **Selected features from PAC-CFC estimator**

|  |  |  |  |  |  |  |  |  |
| --- | --- | --- | --- | --- | --- | --- | --- | --- |
| $\gamma_1$ | ✓ | | ✓ | | | | | |
| $\beta_2$ | | | ✓ | | | | | |
| $\beta_1$ | ✓ | ✓ | ✓ | | ✓ | | | |
| $\alpha_2$ | | | | ✓ | | | | |
| $\alpha_1$ | ✓ | ✓ | | | | | | |
| $\theta$ | ✓ | ✓ | | | | | | |
| high-δ | ✓ | | | | | | | |
| low-δ | | | | | | | | |
| | low-δ | high-δ | $\theta$ | $\alpha_1$ | $\alpha_2$ | $\beta_1$ | $\beta_2$ | $\gamma_1$ |

**Table 2.** **Selected features from AAC-CFC estimator**

|  |  |  |  |  |  |  |  |  |
| --- | --- | --- | --- | --- | --- | --- | --- | --- |
| $\gamma_1$ | | | | | | | ✓ | |
| $\beta_2$ | | | | | ✓ | ✓ | | |
| $\beta_1$ | | | | | | | | |
| $\alpha_2$ | | | | ✓ | | | | |
| $\alpha_1$ | | | ✓ | | | | | |
| $\theta$ | ✓ | ✓ | | | | | | |
| high-δ | ✓ | | | | | | | |
| low-δ | | | | | | | | |
| | low-δ | high-δ | $\theta$ | $\alpha_1$ | $\alpha_2$ | $\beta_1$ | $\beta_2$ | $\gamma_1$ |

**Table 3.** **Selected features from CMI-CFC estimator**

|  |  |  |  |  |  |  |  |  |
| --- | --- | --- | --- | --- | --- | --- | --- | --- |
| $\gamma_1$ | | | | ✓ | ✓ | | | |
| $\beta_2$ | | ✓ | | | ✓ | | | |
| $\beta_1$ | ✓ | | ✓ | | ✓ | | | |
| $\alpha_2$ | | | ✓ | | | | | |
| $\alpha_1$ | | | ✓ | | | | | |
| $\theta$ | | | | | | | | |
| high-δ | | | | | | | | |
| low-δ | | | | | | | | |
| | low-δ | high-δ | $\theta$ | $\alpha_1$ | $\alpha_2$ | $\beta_1$ | $\beta_2$ | $\gamma_1$ |

**Table 4.** **Selected features from MI-CFC estimator**

|  |  |  |  |  |  |  |  |  |
| --- | --- | --- | --- | --- | --- | --- | --- | --- |
| $\gamma_1$ | | | | | | | | |
| $\beta_2$ | | | | | | | | |
| $\beta_1$ | | | | | | | | |
| $\alpha_2$ | | | | | | | | |
| $\alpha_1$ | | | | | | | | |
| $\theta$ | ✓ | ✓ | | | | | | |
| high-δ | ✓ | | | | | | | |
| low-δ | | | | | | | | |
| | low-δ | high-δ | $\theta$ | $\alpha_1$ | $\alpha_2$ | $\beta_1$ | $\beta_2$ | $\gamma_1$ |

**Table 5.** **Selected features from RP.**

|  |  |  |  |  |  |  |  |  |
| --- | --- | --- | --- | --- | --- | --- | --- | --- |
| | low-δ | high-δ | $\theta$ | $\alpha_1$ | $\alpha_2$ | $\beta_1$ | $\beta_2$ | $\gamma_1$ |
| **RP** | ✓ | ✓ | ✓ |  |  |  |  |  |

Fig.6 illustrates the comodulograms of the four CFC estimators adopted in the present study averaged across epochs at each sleep stage from a single-subject. On the x-axis are the frequencies of the modulator while on the y-axis the frequencies of the modulated brain rhythm. Clearly, one can detect the differentiation of the strength of the coupling across sleep stages for PAC (Fig.6A), AAC (Fig.6B) and CMI (Fig.6C) compared to MI (Fig.6D) which mostly detected modulations between low,high δ and θ brain rhythms. PAC coupling was elevated between low, high δ and the rest of frequency subcomponents in deep sleep compared to NREM and W.

Fig.7 illustrates the RP of each frequency subcomponent at each sleep stages and the wake period from a single-subject (same as in Fig.6) after averaging across all epochs from a single-scan. It is obvious that low-δ {0.1-1.5 Hz}, high-δ (K-Complex) {1.6-4 Hz} and θ {4-8 Hz} demonstrated significant differences between the sleep stages and the wake period.

**Fig.6.**
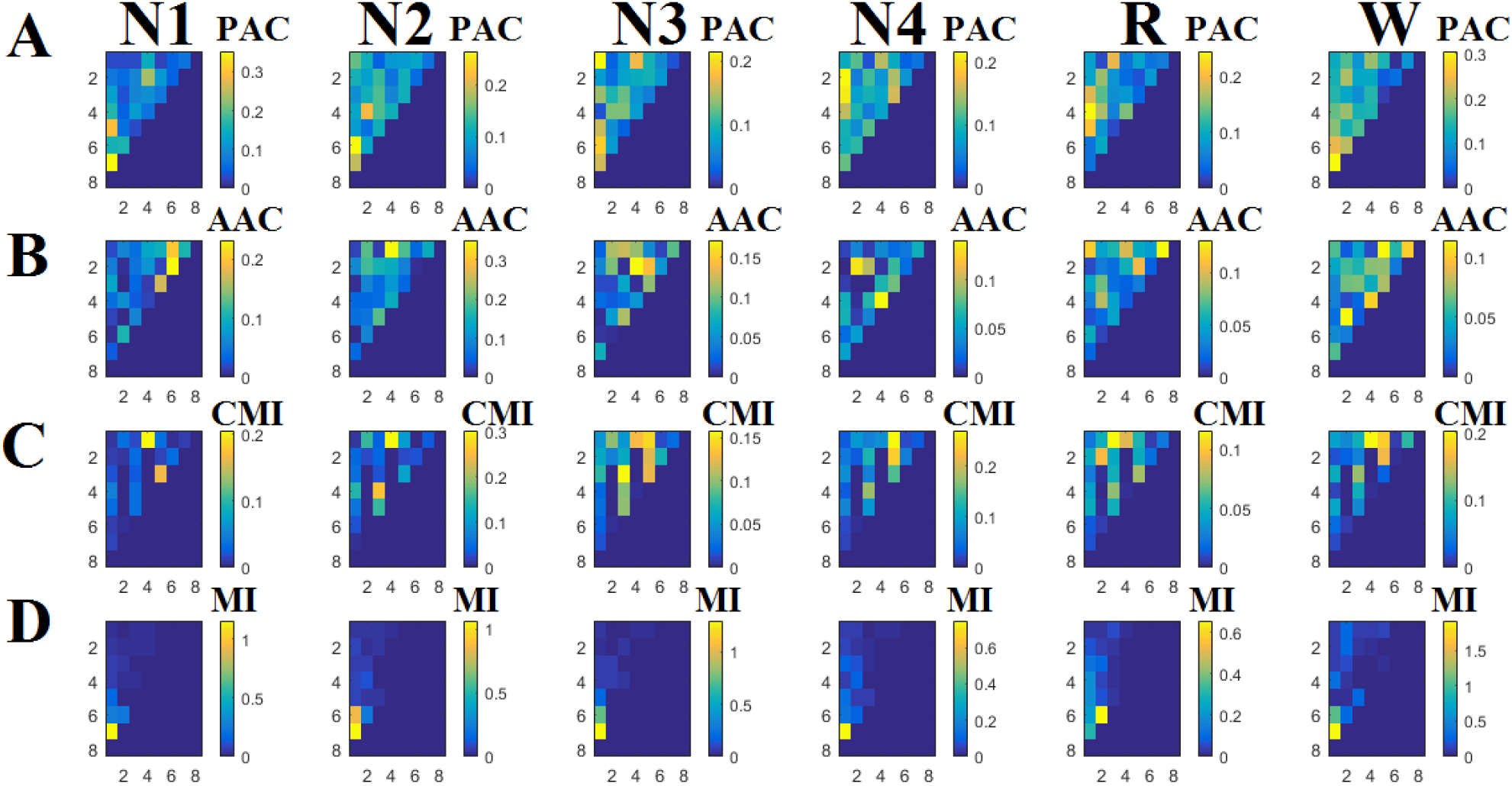
Illustration of the comodulograms of the four CFC estimators adopted in the present study averaged across epochs at each sleep stage from a single-subject. On the x-axis are the frequencies of the modulator while on the y-axis the frequencies of the modulated brain rhythm. (NREM1-4: N1, N2, N3,N4, R:REM,W:WAKE; Frequencies 1-8: 1:low-δ {0.1-1.5 Hz}, 2:high-δ (K-Complex) {1.6-4 Hz}, 3:θ {4-8 Hz},4:α_1_ {8-10 Hz}, 5:α_2_ {10-13 Hz},6:β_1_ (spindle) {14-20 Hz},7:β_2_ {21-30 Hz} and 8:γ_1_ {31 – 45 Hz}).

**Fig.7.**
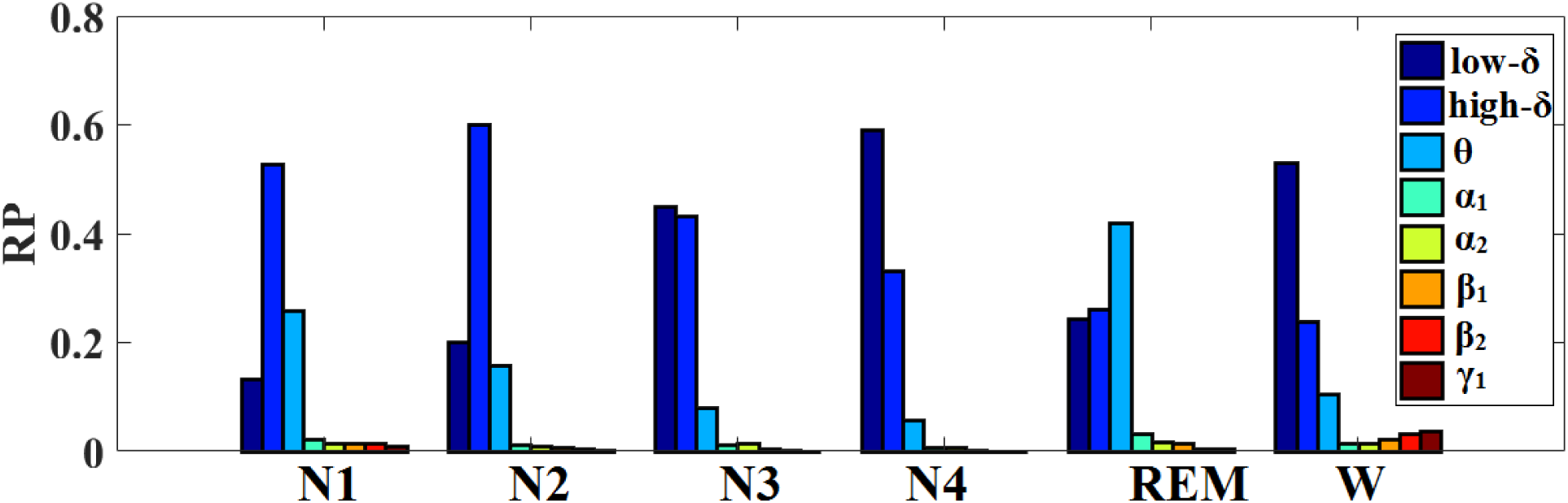
Illustration of the relative power (RP) averaged across epochs at each sleep stage from a single-subject. On the x-axis are presented in six blocks the RP of eight frequency bands. (NREM1-4: N1, N2,N3,N4, R:REM,W:WAKE; Frequencies 1-8: 1:low-δ {0.1-1.5 Hz}, 2:high-δ (K-Complex) {1.6-4 Hz}, 3:θ {4-8 Hz},4:α_1_ {8-10 Hz}, 5:α_2_ {10-13 Hz},6:β_1_ (spindle) {14-20 Hz},7:β_2_ {21-30 Hz} and 8:γ_1_ {31 – 45 Hz}).

### 3.2 Sleep State Classification Performance

We achieved very high classification accuracy, sensitivity and specificity of **96.2 ± 2.2%, 94.2 ± 2.3%**, and **94.4 ± 2.2% respectively across the 20 folds**. Complementary, mean F1-score was also high (92%, range 90–94%). The aforementioned results were succeeded with the application of multi-class Bayes Naive classifier (with kernel). All our measures were first averaged across folds at each repetition and then we estimated the mean and standard deviation across 100 repetitions of the 20-fold.

The best results were achieved with the Fpz-Cz sensor. Table 6 demonstrates the averaged confusion matrix across the 20 folds. Diagonal elements referred to the matching in % between experts and our algorithm while off-diagonal elements illustrate the % of mismatch. The most correctly classified sleep stage was the N4 following by N3, W, N2, R and lastly the N1. Most misclassifications of N1 were happened in R and W stage, for N2 in N3 and R, for N3 in R and W, for N4 in N3, for R in NI and N3 and for W in N4 and R.

**Table 6.** Averaged Confusion Matrix Across the 20 Folds based on the best performance Fpz-Cz channel (%).

|  | N1<br>(Algorithm) | N2<br>(Algorithm) | N3<br>(Algorithm) | N4<br>(Algorithm) | R<br>(Algorithm) | W<br>(Algorithm) |
| --- | --- | --- | --- | --- | --- | --- |
| N1 (Expert) | 94.1<br>(4.9) |  |  |  | 0.6<br>(0.03) |  |
| N2 (Expert) |  | 94.7<br>(4.1) |  |  |  |  |
| N3 (Expert) | 0.03<br>(0.02) | 2.8<br>(0.03) | 98.8<br>(3.8) | 0.09<br>(0.03) | 5.1<br>(0.06) |  |
| N4 (Expert) |  |  |  | 99.2<br>(3.7) |  | 3.7<br>(0.06) |
| R (Expert) | 3.2<br>(0.03) | 3.1<br>(0.05) | 0.07<br>(0.02) |  | 94.6<br>(4.1) | 0.8<br>(0.02) |
| W (Expert) | 3.5<br>(0.04) |  | 0.06<br>(0.02) |  |  | 95.9<br>(4.7) |

All our measures (F1 score, accuracy, sensitivity, specificity) that evaluated the robustness of the proposed scheme for the automatic sleep stage scoring based on a single-sensor are superior to previous attempts on the same dataset (Berthomier et al., 2007; Liang et al., 2012; Tsinalis et al., 2016). Complementarily, our methodology worked independently for N3 and N4 without merging them into a single stage.

Figure 8 illustrates the manually scored hypnogram of a subject versus the proposed single-sensor automatic sleep stage classification.

**Figure 8.**
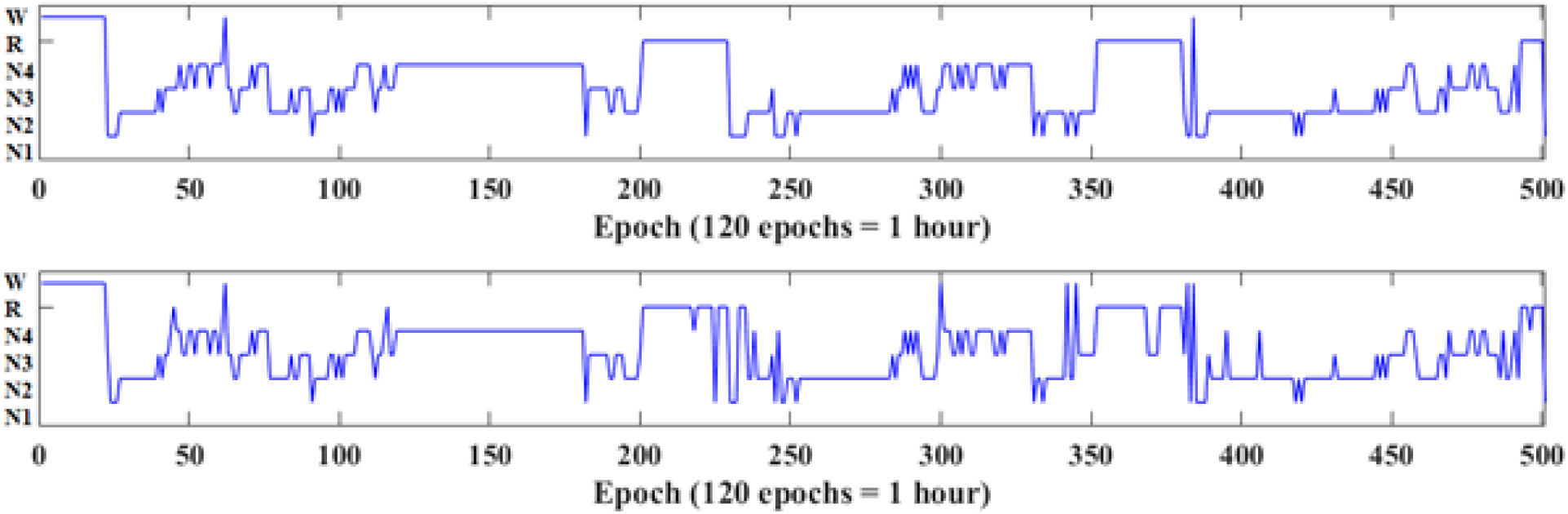
**Manual scoring versus automatic sleep stage scoring.** On the top, we illustrated the original manually scored hypnogram while in the bottom the estimated hypnogram using the proposed algorithm for the second night of subject 1.

### 3.3 Automatic Sleep Stage Claassification Performance (ASSC) in an External Dataset

We achieved very high classification accuracy, sensitivity and specificity also for the second external dataset. Specifically, we achieved **95.1 ± 2.9%, 94.1 ± 3.1%**, and **94.0 ± 2.9% respectively across the subjects of the second dataset**. Complementary, mean F1-score was also high (91%, range 89–92%). The aforementioned results were succeeded with the application of multi-class Bayes Naive classifier (with kernel) and using as a training set the features extracted from the first dataset. All our measures were averaged across subjects.

Table 7 demonstrates the averaged confusion matrix across the 77 subjects. Diagonal elements demonstrate the matching in % between expert and our algorithm while off-diagonal elements illustrate the % of mismatch. The most correctly classified sleep stage was the N4 following by N3,W,N2, R and lastly the N1. Most misclassifications of N1 were happened in R and W stage, for N2 in N3 and R, for N3 in R and W, for N4 in N3, for R in NI and N3 and for W in N4 and R.

**Table 7.** Averaged Confusion Matrix Across the 77 subjects based on the best performance Fpz-Cz channel (%).

|  | N1<br>(Algorithm) | N2<br>(Algorithm) | N3<br>(Algorithm) | N4<br>(Algorithm) | R<br>(Algorithm) | W<br>(Algorithm) |
| --- | --- | --- | --- | --- | --- | --- |
| N1 (Expert) | **94.5**<br>(4.9) |  |  |  | 1.1<br>(0.05) |  |
| N2 (Expert) |  | **94.2**<br>(4.1) |  |  |  |  |
| N3 (Expert) | 0.03<br>(0.04) | 2.9<br>(0.05) | **97.6**<br>(3.8) | 3.4<br>(0.04) | 5.3<br>(0.07) |  |
| N4 (Expert) |  |  |  | **96.8**<br>(3.7) |  | 3.5<br>(0.07) |
| R (Expert) | 3.0<br>(0.05) | 3.2<br>(0.04) | 2.1<br>(0.02) |  | **94.1**<br>(4.1) | 1.7<br>(0.05) |
| W (Expert) | 3.6<br>(0.04) |  | 1.8<br>(0.02) |  |  | **95.1**<br>(4.7) |

### 3.4 Relative Signal Power Changes during Sleep

Based on our results of relative signal power changes across the sleep stages (Fig.8), low-δ {0.1-1.5 Hz} is elevated during deep sleep starting from N1 to N4. Signal power of high-δ (K-Complex) is higher in N1 and N2 and is decreased till REM. Complementary, the signal power of θ {4-8 Hz} is higher in REM which is an indicator of the activity of the brain during REM stage. We didn’t detect significant relative power changes in β_1_ frequency across the sleep stages and especially in N2 linked to sleep spindles maybe because of the long epoch of 5 sec.

### 3.5 Cross-Frequency Coupling Changes during Sleep

By combining Tables 1-4 with Fig.7, we revealed important tendencies of increased or decreased patterns of CFC with the four adopted estimators during the progression to deep sleep. For PAC estimator, we have detected higher values for N3-N4 sleep stages especially for slower brain rhythms (low-δ {0.1-1.5 Hz}, high-δ {1.6-4 Hz}, θ) while during REM, higher values were detected between θ-γ_1_, α_1_-α_2_ and α_2_-β_1_. For AAC estimator, the coupling for frequency pairs low-δ - high-δ, low-δ – θ and high-δ– θ is elevated during N3-N4, θ-α_1_ is higher in N3 stage, α_1_- α_2_ and α_2_-β_2_ are higher in N4 while β_2_-γ_1_ is higher in REM. Complementary, β_1_-β_2_ interactions were higher in N1, decreased in N2 and preserved stable during N3-N4 and REM. The complex version of MI (CMI) works better with θ and α_2_ brain rhythms that further complements PAC estimates. Specifically, we observed higher values for low-δ-β_1_ and high-δ-β_2_ in N3 stage while θ-α_1_ increased during deep sleep starting from N2 up to N4. On the opposite, θ- α_2_ is higher in N2 and is decreased till N4. θ- β_1_ demonstrates higher values for N3 and further decreased in N4 and REM stages. α_1_- γ_1_ demonstrates higher values in N1-N2 and then decreased till REM. α_2_- β_1_,α_2_- β_2_ and α_2_- γ_1_ increased during the first three sleep stages reaching the maximum in N3 and then further decreased till REM. Finally, the MI contributes to the multiparametric classifier with three features. low-δ- high-δ demonstrates high values in the N1-N2-N3 sleep stages and further reduced till the REM sleep. low-δ- θ increased from N1 to N2 and then reduced progressively till REM sleep. Finally, high-δ- θ is one of the detectable cross-frequency couplings in N1-N2 with MI with very low values in N3 and higher values in REM.

## 4 Discussion

In the present study, we demonstrated the feasibility of a novel single-sensor automatic sleep stage classification (ASSC) algorithm based on different aspects of cross-frequency coupling estimates. We achieved a very high classification sensitivity, specificity and accuracy of **90.3 ± 2.1%, 94.2 ± 2.3%**, and **94.4 ± 2.2% across 20 folds**, respectively with high mean F1-score (92%, range 90–94%) when multi-class Bayes Naive classifier (with kernel). Our results revealed that Fpz-Cz sensor is the most appropriate for ASSC succeeding better performance compared to Pz-Oz. Complementary, we replicated our excellent results in an external second open sleep database of 77 subjects. Specifically, we achieved very high classification accuracy, sensitivity and specificity namely **95.1 ± 2.9%, 94.1 ± 3.1%**, and **94.0 ± 2.9% respectively across the subjects of the second dataset**. Additionally, mean F1-score was also high (91%, range 89–92%).

The whole methodology and the novelty of the proposed scheme can be summarized as follow:

- We analysed EEG recordings at every 5 s instead of 30 s epochs (which is the epoch length for the expert scoring)
- We kept N3 and N4 as single sleep stages without attempting to merge them (Tsinalis et al., 2016)
- We adopted MODWT to simultaneously decompose the EEG recordings into true activity assigned to one of the basic frequency ranges and also to denoise the EEG recordings from eye-movements and muscle activity
- We adopted apart from relative signal power, four different connectivity estimators (three phase-to-amplitude: PAC,MI,CMI and one amplitude-to-amplitude: AAC)
- We used a dataset where all the subjects (expect one) had more than one recording
- Cross-validation scheme was designed such as to:

a. To access the generalisability of the method by adopting a 20-fold where at each fold only the recording from a single subject were used for training while each subjects’s recording were used only once for testing
b. Avoid imbalanced sleep stages distribution on the training set. For that purpose, we repeated the 20-fold CV, 100 times by randomly sampling the class labels across sleep stages and subjects such as to keep their representation equal.
- Our results outperformed previous comparable studies on the same dataset (Berthomier et al., 2007; Liang et al., 2012; Tsinalis et al., 2016)
- We also replicated the proposed ASSC method based on CFC features in a second sleep dataset
- We demonstrated for the first the effectiveness of different cross-frequency coupling estimates to the improvement of CFC and also a better understanding of the multiplexity of the human brain in sleep

Our methodology presents a single-sensor automatic sleep stage classification based on cross-frequency coupling estimates that succeeded a very high classification performance. CFC outperformed trivial features derived from sleep recordings supporting further the notion of CFC for the sleep stage classification. A previous study attempted to demonstrate the positive effect of CFC on ASSC succeeding an overall accuracy of 75 % with four sleep stages (Sanders et al., 2014). A more recent study explored the phase-to-amplitude coupling in deep sleep and in epileptic patients where they observed elevated PAC (Amiril et al., 2015).

Brain rhythms can interact between each others with different mechanisms like phase-to-phase, phase-to-amplitude (PAC) and amplitude-to-amplitude envelope correlation (see Figure 2 in Buszaki et al., 2012). An important mechanism that exists in a typical scenario when two oscillators with the same or different frequency within the same or different anatomical brain area are entrained each other is phase-to-phase coupling. This is a mechanism that mainly can be estimated between a pair of sensors and not within a sensor (Dimitriadis et al., 2015b). A less temporally less precise, but nevertheless important, brain interaction between oscillators of similar or different frequency is expressed by the temporal covariation of their amplitude/power, known as amplitude/power comodulation or amplitude-amplitude/power-power coupling (correlation of the envelope: Bruns and Eckhorn,2004).

The third basic mechanism for brain interactions is called frequency phase-to-amplitude (CPAC or PAC; Pittman-Polletta et al., 2014; Dimitriadis et al., 2015a, 2016a,b) coupling of nested oscillations. One reason why slow oscillations couple faster brain rhythms in multiple brain areas is explained with the conduction velocities of cortical neurons. Slower oscillators compared to faster activate more neurons in a larger volume (von Stein and Sarnthein, 2000) and are associated with larger membrane potential changes since in longer temporal windows, a large portion of spikes of many more upstream neurons can be integrated (Hasenstaub et al., 2005; Quilichini et al., 2010).

PAC has been reported between every pair of brain rhythms in interactive circuits in the mammalian cortex from low oscillations as 0.025 Hz up to fast as 500 Hz (Sirota et al., 2003). For example, the occurrence of hippocampal “ripples” (40 to 200 Hz) is coupled to dendritic layer sharp waves and phase-modulated by sleep spindles (12 to 16 Hz). In turn, the spindle-modulated sharp wave-ripple complex is phase-coupled to neocortical slow oscillations (0.5 to 1.5 Hz) (Sterlade et al., 1993; von Stein and Sarnthein, 2000; Sirota and Buzsaki, 2005; Isomura et al., 2006; Ji and Wilson,2007;Peyrache,2011) and all these rhythms are modulated by the ultraslow (0.1 Hz) oscillation (Sirota et al., 2003). Finally, the hierarchy of brain oscillators is formed by the CFC interactions that further support the interaction of multiple brain rhythms across spatial and temporal scales (Buzsaki et al., 2012). Here, using a single EEG sensor, we have detected higher values for N3-N4 sleep stages especially for slower brain rhythms (low-δ {0.1-1.5 Hz}, high-δ {1.6-4 Hz}, θ) while during REM stage, higher values were detected between θ- γ_1_, α_1_-α_2_ and α_2_- β_1_.

The formation of new memories demands the coordination of neural activity across widespread brain regions. In both humans and animals, the hippocampus is believed to support the formation of new associative or contextually mediated memories (Clemens et al., 2009). During the consolidation of new memories on a system-level, mnemonic representations of items initially reliant on the hippocampus and after are thought to travel to neocortical sites for more permanent storage. Sleep has this privilege role for facilitating this information transfer (Born and Wilhelm,2012). Mechanistically, consolidation processes have been proved to be rely on systematic interactions between the three basic neuronal oscillations that characterizing non–rapid eye movement (NREM) sleep: slow-oscillations, spindles and ripples (Staresina et al.,2015). Staresina et al., (2015) used direct intracranial EEG recordings from human epilepsy patients during natural sleep to test the assumption that slow-oscillations, spindles and ripples are functionally coupled with the hippocampal activity. They demonstrated a PAC between 0.5–1.25 Hz (slow-oscillation range) and 12–16 Hz (spindle range), respectively using EEG modality and CZ sensor. In addition, they demonstrated a hippocampal PAC during NREM sleep stages providing also a link between EEG-CZ recordings with hippocampal estimates from epileptic patients. The hierarchical role of these three sleep components (slow-oscillations, spindles and ripples) was revealed via phase-to-amplitude coupling based on mean vector length estimator.

In the present study, we revealed multiple cross-frequency interactions between slow, medium frequencies and low γ activity in NREM sleep with every cross-frequency coupling estimator. Our results further support that CFC during NREM sleep are significant attributes of sleep and the overall memory consolidation. The β_1_- β_2_ interactions revealed with AAC should be further explored in direct link with their functionality in the consolidation of memories.

A recent study untangled hippocampo-cortical CFC as the basic mechanisms mediates memory consolidation during sleep (Maingret et al., 2016). They provided a clear functional link between sharp-wave ripples, delta waves and ripples. Logothetis et al., (2012) demonstrates the CFC-PAC coupling between hippocampo-cortical areas during a subcortical silence and off-line memory consolidation while Amiri et al., (2016) demonstrates an enhanced PAC in deep sleep and also in the onset zone of focal epilepsy.

In the present study, we demonstrated for the first time the effectiveness of different aspects of CFC namely, PAC and amplitude-to-amplitude envelope correlation to automatically classify sleep stages (Tables 1-4 and Fig.6). Additionally, we analysed sleep data under CFC for the first time in healthy populations further proved that CFC interactions exist during sleep showing and that were altered during the transition between sleep stages. We succeeded high classification accuracy in two datasets (training and testing dataset) using the activity recorded from a single EEG sensor. In the second study, we achieved lower mean classification accuracy across sleep stages compared to the first one and especially in N4 and R sleep stages (Table 6 vs Table 7). One possible explanation on these findings could be the amplitude differences of slow oscillations between gender and across the lifespan (Mourtazaev et al., 1995). It is well-known that aging affects the neurophysiological generation of slow-wave oscillations (Leirer et al., 2011).

Here, we selected the relative power of low-δ {0.1-1.5 Hz}, high-δ (K-Complex) {1.6-4 Hz} and θ {4-8 Hz} as complementary features to the CFC estimates for a better classification of the sleep stages (Table 5 and Fig.7). δ waves were defined within the range of 1-4 <u>Hz</u> (Walker,1990). Compared to the others brain waves, δ waves have the highest amplitude while recent studies described slower (<0.1 Hz) oscillations (Hiltunen et al., 2014). Both sleep stages 3 and 4 are dominated by δ waves with higher representation in sleep stage 4 (Iber et al., 2007). In addition, δ waves are often associated with another EEG phenomenon, the <u>K-complex</u> (high-δ: {1.6-4 Hz}). K-Complexes have been demonstrated to precede δ waves in slow wave sleep (De Gennaro et al., 2000). Both K-complex and δ <u>wave</u> activity in sleep stage 2 generate both slow-wave (∼0.8 Hz) and δ (1.6–4.0 Hz) oscillations. However, their topographical distribution is different while the δ power of the K-complexes is higher (Happe et al., 2002). Additionally, δ waves can be further classified according to the location where they mostly detected into: frontal (FIRDA), temporal (TIRDA), and occipital (OIRDA) intermittent δ activity (Brigo,2011).During sleep, θ activity appears as the prominent EEG activity in REM sleep while θ activity serves as the background activity of both spindles and K-complexes during sleep stage 2 (Rodenbeck et al., 2006).

Various CFC-PAC estimates and AAC have been employed here to estimate the alternative CFC mechanisms of brain’s communication between different frequencies. Fig.6 illustrates the comodulograms of the four CFC estimators adopted in the present study averaged across epochs at each sleep stage from a single-subject. Clearly, one can detect the differentiation of the strength of the coupling across sleep stages for PAC (Fig.6A), AAC (Fig.6B) and CMI (Fig.6C) compared to MI (Fig.6D) which mostly detected modulations between low,high δ and θ brain rhythms. PAC coupling was elevated between low,high δ and the rest of frequency subcomponents in deep sleep compared to NREM and W. Our observations are direct linked with the notion that many frequencies are modulated by the ultraslow (0.1 Hz) oscillation during sleep (Sirota et al., 2003) affected by different sleep stages (Staresina et al.,2015).Enhanced PAC during deep sleep was also observed in epileptic patients (Arimi et al., 2016).

Our main goal was to further enhance the efforts of succeeding ASSC with a high accuracy from a single-sensor. It is also significant to study simultaneously intra and inter-frequency interactions including phase-to-phase between multi EEG sensors in order for a better understanding of the mechanism and interactions occurred during sleep. It is important to mention here that our results were externally validated with a second big dataset which makes the whole analysis stronger and unbiased from the subjective sleep scoring based on which we trained our classifiers in the first dataset.Our next goal is to use an open database and to extend present and also our previous work where we explored intra-frequency interactions between multiple EEG sensors during the five sleep stages (Dimitriadis et al., 2009).

Apart from the studying of CFC interactions within a single-sensor, the whole analysis based on a healthy population. For that reason, our next goal is to explore brain connectivity during sleep stages using more EEG sensors and all the possible intra and inter-frequency coupling modes. For that reason, we will work on ISRUC database (multi-channels; Sirvan et al., 2016) which provides a large dataset with healthy individuals and also sleep recordings from non-healthy individuals.

To the best of our knowledge, our method achieved the best performance in the literature when classification is done across all five sleep stages and wake (six class) condition simultaneously using a single channel of EEG. This is different from adopting a one versus all classification strategy where the classification performance is over-estimated. Our results outperformed also studies that they used more than one channel (EOG,EEG,EMG) to extract features for the accurate sleep scoring. Our next goal is to test our algorithm on low-cost commercial EEG sensors (ear-EEG; Looney et al., 2016) and on recordings from home environments and to further explore CFC patterns in sleep disorders.

## Acknowledgements

SID and DL were supported by MRC grant MR/K004360/1 (Behavioural and Neurophysiological Effects of Schizophrenia Risk Genes: A Multi-locus, Pathway Based Approach). SID is also supported by MARIE-CURIE EU-UK COFUND FELLOWSHIP.

